## Supplementary Fig. 1 for "A rendezvous of two second messengers: The c-di-AMP receptor protein DarB controls (p)ppGpp synthesis in *Bacillus subtilis*"

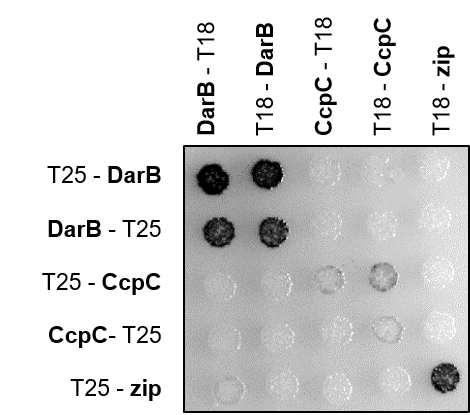

**Supplementary Fig. 1. DarB does not interact with CcpC.** Bacterial two-hybrid (BACTH) experiment testing for the interaction of DarB with CcpC. N- and C-terminal fusions of DarB and the CcpC variants to the T18 or T25 domain of the adenylate cyclase (CyaA) were created and the proteins were tested for interaction in *E. coli* BTH101. Dark colonies indicate an interaction that results in adenylate cyclase activity and subsequent expression of the reporter β-galactosidase. While DarB exhibits self-interaction, no interaction between DarB and CcpC could be detected.

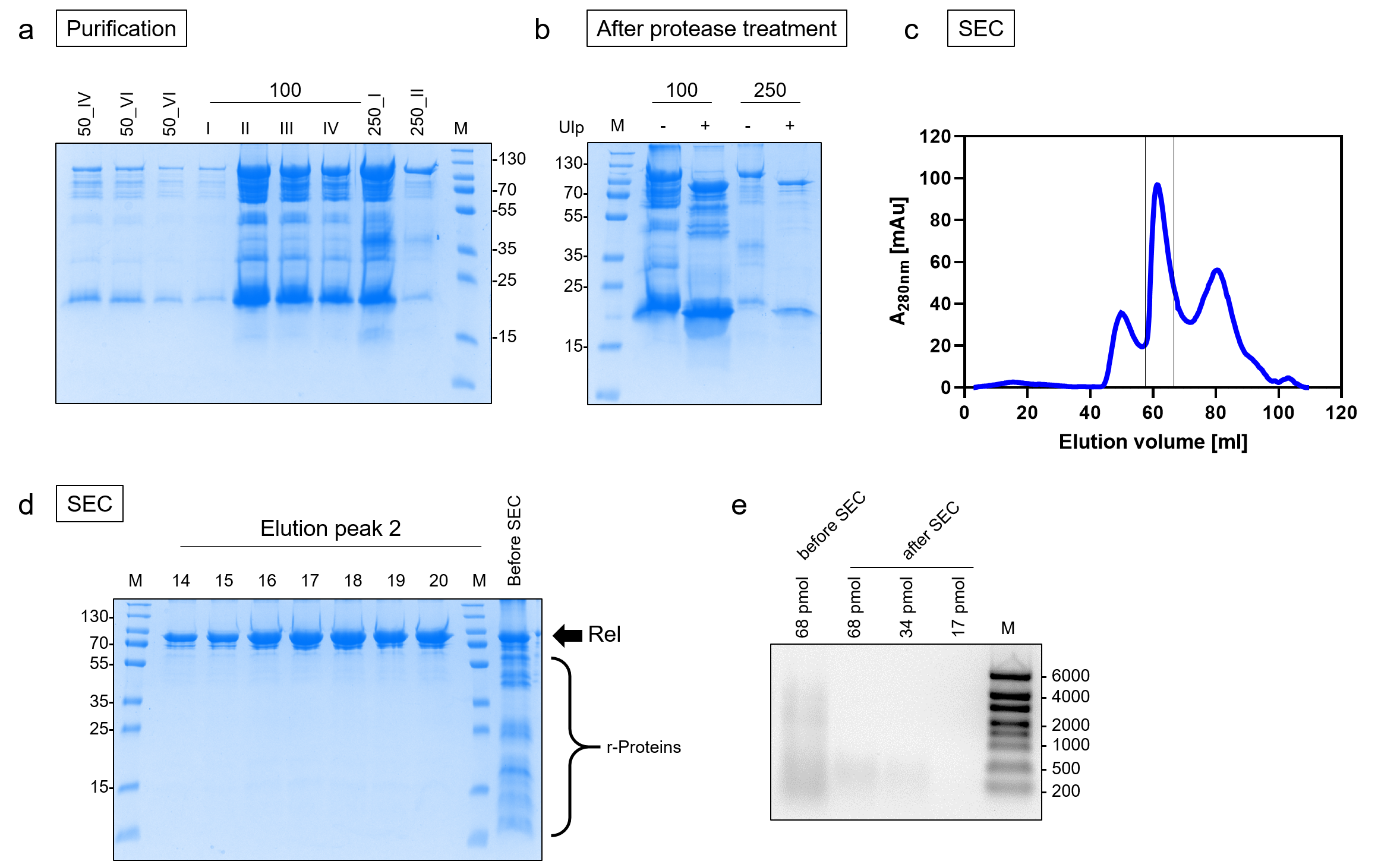

**Supplementary Fig. 2. Purification of Rel. a**, 10xHis-SUMO-tagged Rel (pVHP186) was overexpressed in *E. coli* Rosetta DE3 and purified in 750 mM KCl, 5 mM MgCl_2_, 40 µM MnCl_2_, 40 µM Zn(OAc)_2_, 20 mM imidazole, 10% glycerol, 4 mM β-mercaptoethanol, 25 mM HEPES:KOH pH 8 via a Ni^2+^nitrilotriacetic acid column. The protein was eluted with 100 and 250 mM imidazole and the elution fractions were analyzed by SDS-PAGE and Coomassie staining. **b**, The elution fractions were pooled and the 10xHis-SUMO-tag was cut off by overnight incubation with the SUMO protease. **c**, Size exlusion chromatography (SEC) was performed to remove contamination of the protein preparation. The fractions that were used for the further experiments are highlighted and were analyzed by SDS-PAGE (**d**) and on a denaturing agarose gel to verify the absence of potentially contaminating RNA (**e**). The ratio between A_260_/A_280_ was 0.9 indicating that the protein was almost free of contaminating RNA.

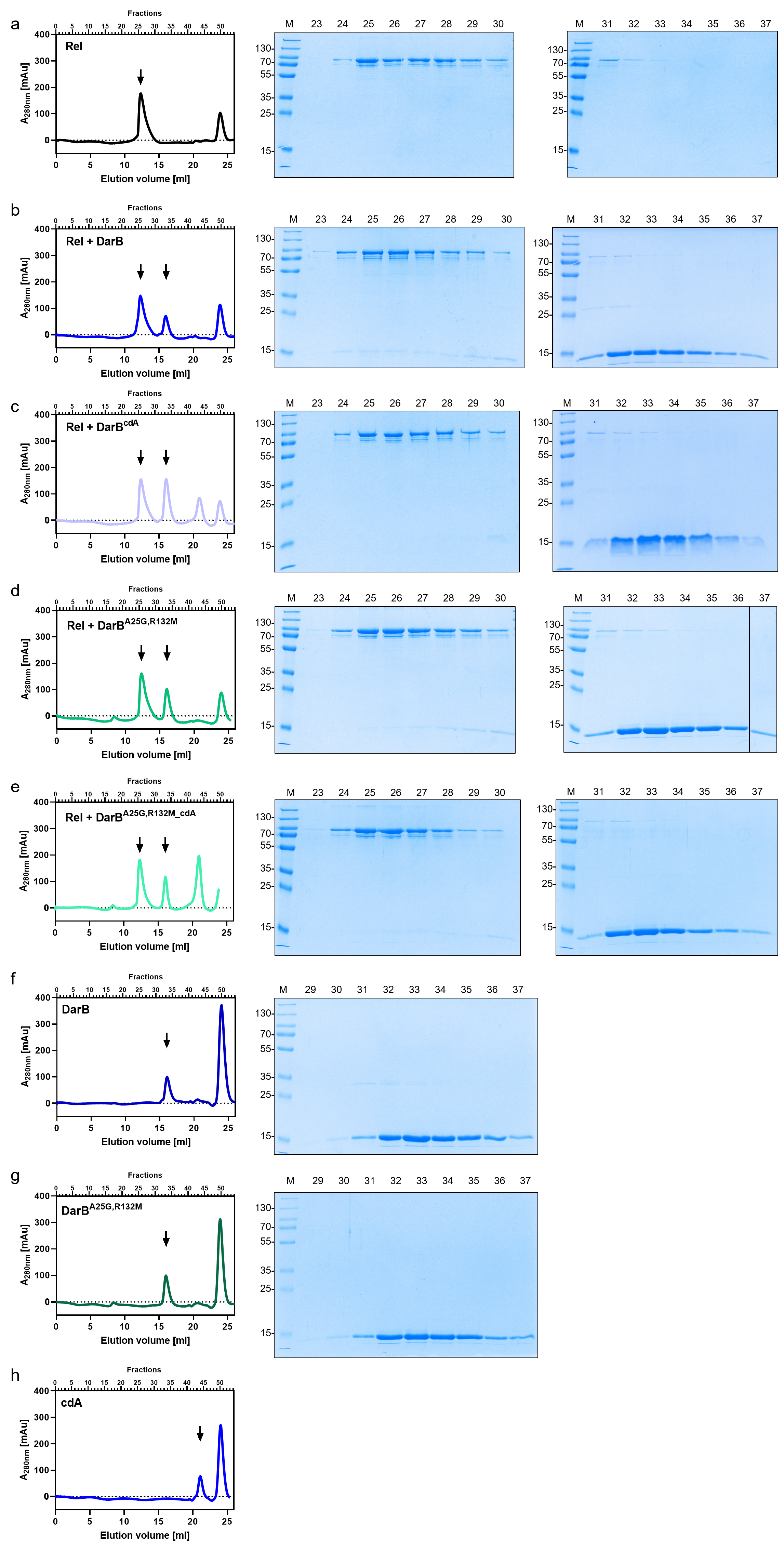

**Supplementary Fig. 3. Analysis of the DarB-Rel-complex by size exclusion chromatography (SEC).** Chromatograms of the SEC run are shown together with the SDS-gels of the relevant elution fractions (indicated above chromatogram and gel). Rel and DarB were used in equimolar concentrations. SEC runs of Rel alone (a) with DarB (b), DarB^cdA^ (c), DarB^A25G,R132M^ (d), or DarB^A25G,R132M_cdA^ (e), and as controls DarB (f), DarB^A25G,R132M^ (g), and c-di-AMP (h). Abbreviation: cdA, c-di-AMP.

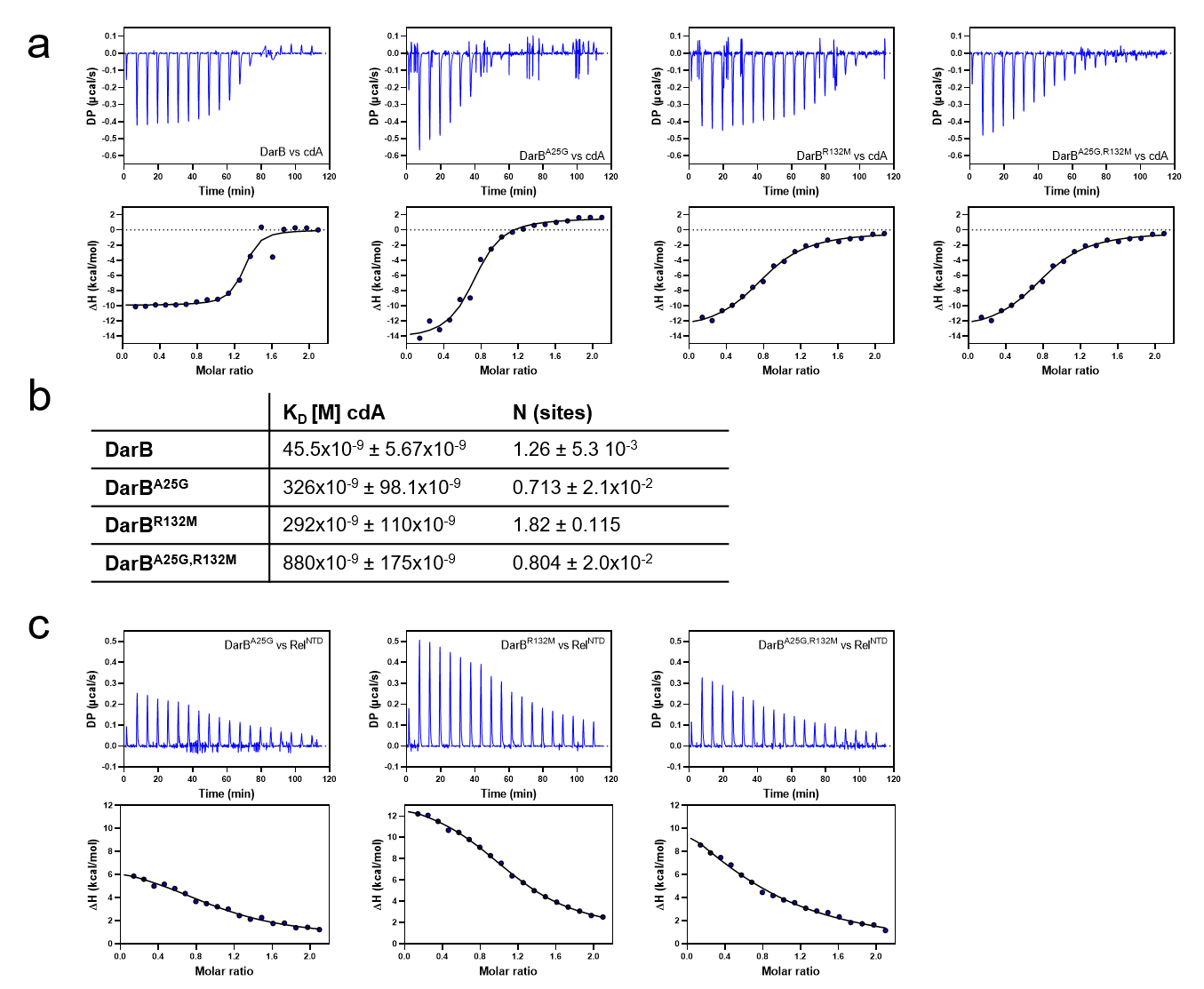

**Supplementary Fig. 4. Analysis of the binding affinities of DarB, and the DarB mutants towards c-di-AMP and Rel^NTD^. a**, The ability of the DarB wild type and the mutant proteins to bind c-di-AMP was assessed by Isothermal titration calorimetry (ITC). The cell and the syringe contained 10 µM DarB and 100 µM c-di-AMP, respectively. Titration profiles and the determined molar ratios. **b**, Calculated K_D_ values for binding of c-di-AMP, as well as the determined number of binding ligand sites. **c**, The interaction of the DarB mutant proteins with Rel^NTD^ was investigated with ITC. The cell and the syringe contained 10 µM Rel and 100 µM DarB, respectively. Abbreviation: cdA, c-di-AMP.

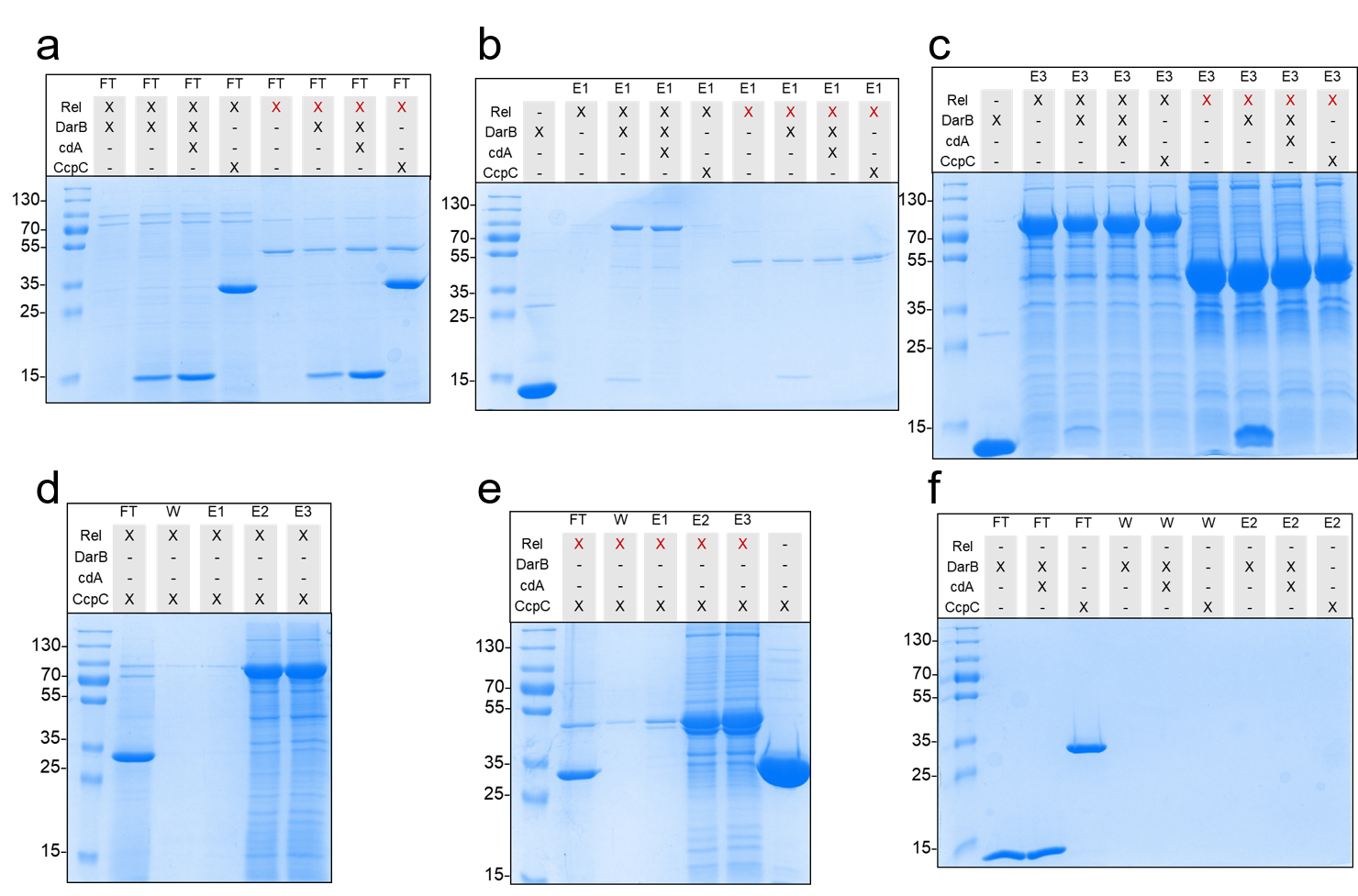

**Supplementary Fig. 5. *In vitro* interaction experiment between DarB and Strep-Rel or Strep-Rel^NTD^.** Strep-Rel or Strep-Rel^NTD^ were immobilized onto a StrepTactin column and incubated with DarB, DarB preincubated with c-di-AMP, or the control protein CcpC. The eluates were analyzed by SDS-PAGE. Control gels are shown, the presence of the N-terminal Strep- Rel^NTD^ variant is indicated by a red cross. **a**, Flow through (FT) after loading of DarB or CcpC; **b**, Elution fractions 1; **c**, Elution fractions 3; **d**, SDS PAGE showing all fractions of the negative control CcpC and Strep-Rel; **e**, SDS PAGE showing all fractions of the negative control CcpC and Strep- Rel^NTD^; **f**, SDS PAGE with all fractions of DarB/ CcpC to exclude unspecific binding to the column. Abbreviation: cdA, c-di-AMP.

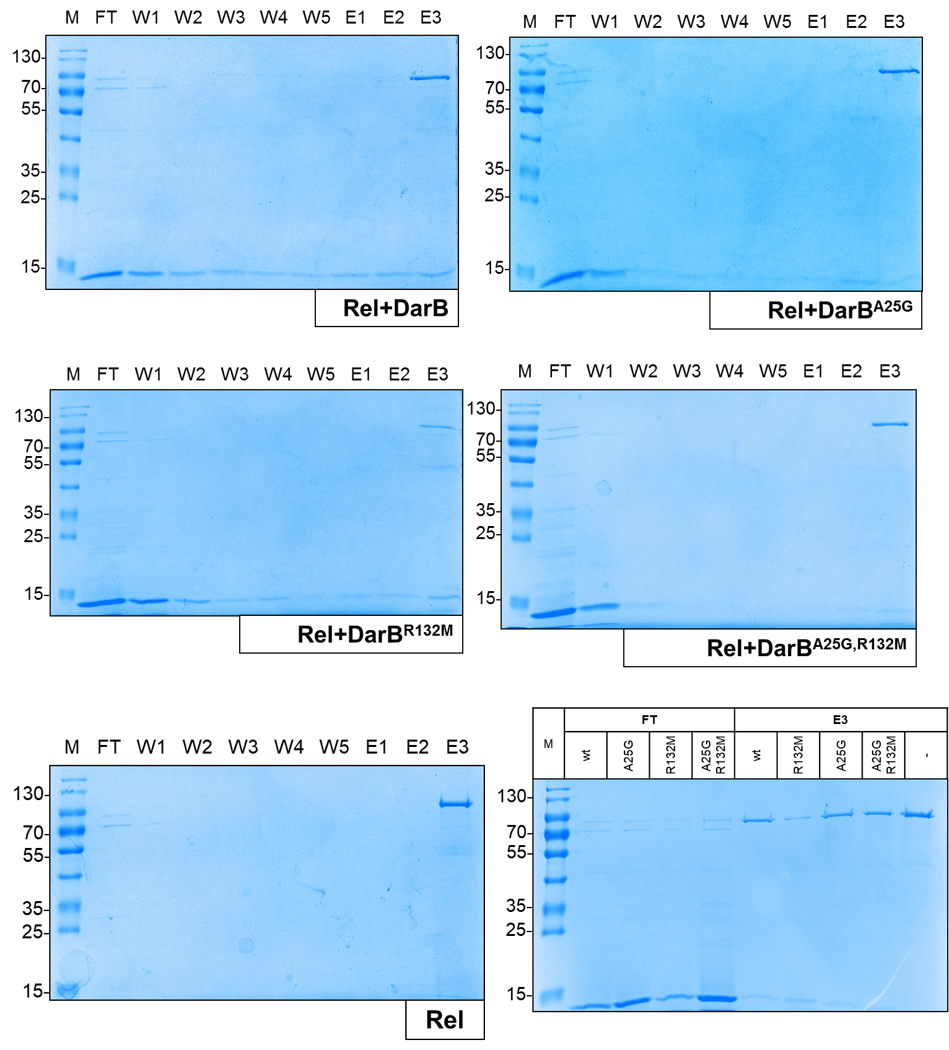

**Supplementary Fig. 6. *In vitro* interaction experiment between DarB, DarB^A25G^, DarB^R132M^, DarB^A25G,R132M^, and Strep-Rel.** Strep-Rel was immobilized onto a StrepTactin column and incubated with DarB, or the DarB mutants. The flow through (FT), the wash fractions (W), and the eluates were analyzed by SDS-PAGE.

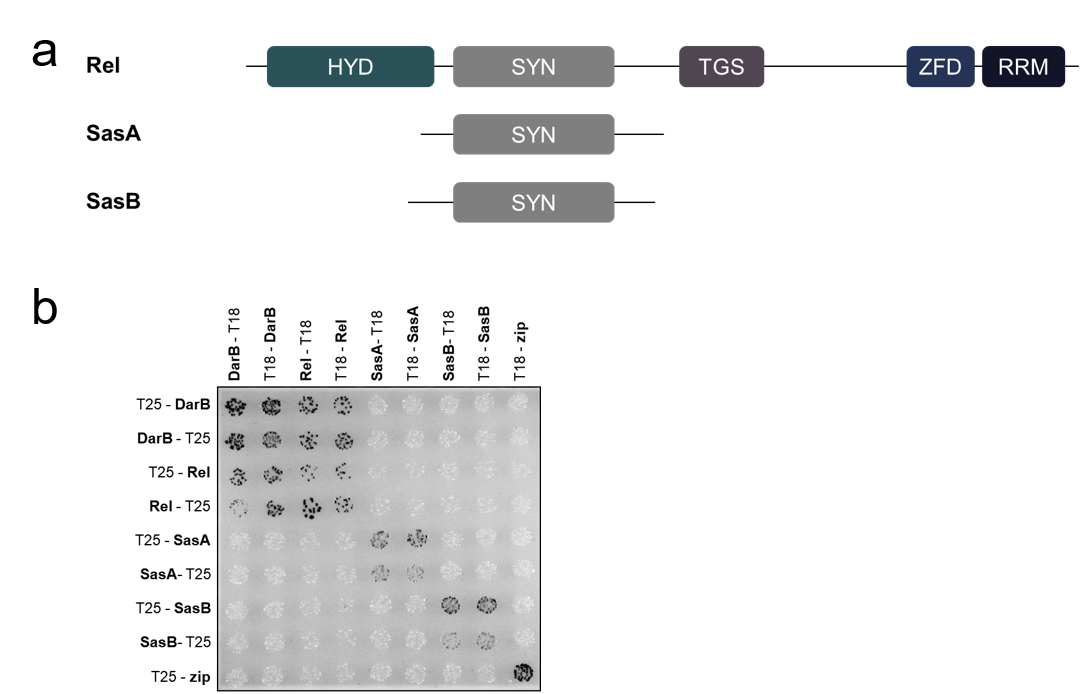

**Supplementary Fig. 7. DarB does not interact with the small alarmone synthetases SasA and SasB.** **a**, Domain organization of Rel and the small alarmone sythetases SasA and SasB. **b**, Bacterial two-hybrid (BACTH) experiment testing for the interaction of DarB with Rel, SasA and SasB. N- and C-terminal fusions of DarB and Rel, SasA or SasB to the T18 or T25 domain of the adenylate cyclase (CyaA) were created and the proteins were tested for interaction in *E. coli* BTH101. Dark colonies indicate an interaction that results in adenylate cyclase activity and subsequent expression of the reporter β-galactosidase. While DarB exhibits an interaction with Rel, no interaction between DarB and SasA or SasB could be detected.

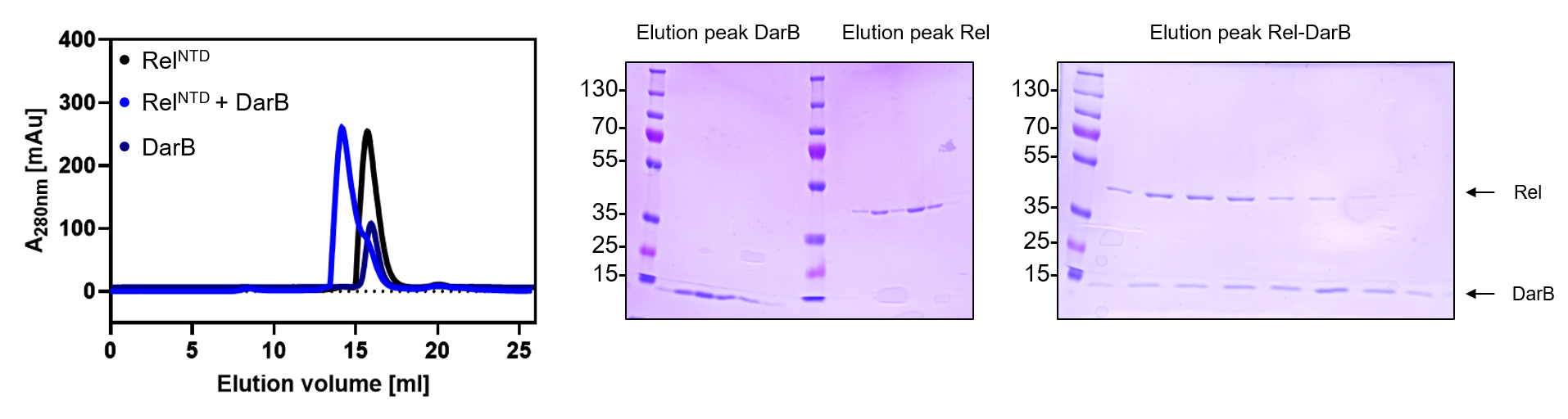

**Supplementary Fig. 8. *In vitro* analysis of the DarB-Rel^NTD^ complex.** Size-exclusion chromatography and multi-angle light scattering (SEC-MALS) of DarB (dark blue), Rel (black), and the DarB-Rel^NTD^-complex (blue). Coomassie-stained gels of the relevant elution fractions are depicted next to the chromatogram.

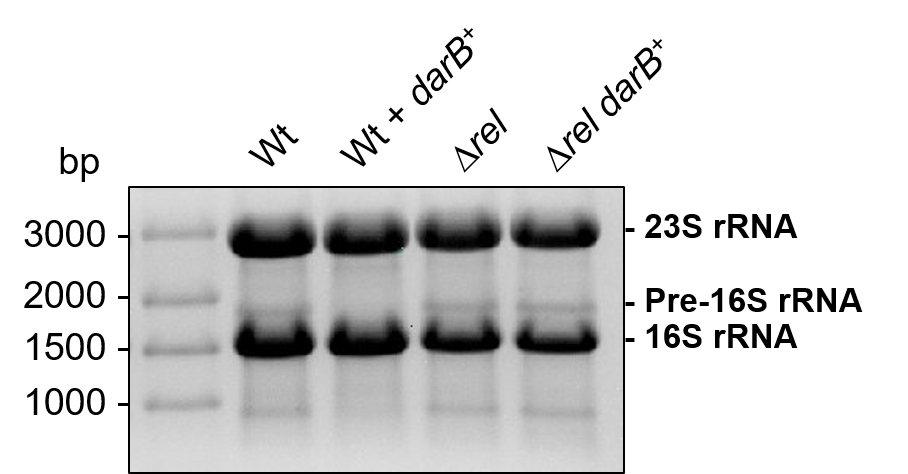

**Supplementary Fig. 9. Analysis of rRNA profiles in *B. subtilis* mutants.** Cultures were grown in minimal medium with 0.1 mM KCl. Total RNA was extracted and 3 µg were resolved on a 1% agarose formaldehyde gel and stained with ethidium bromide. Positions of the ribosomal rRNAs are indicated.

**Supplementary Table 1**

**Analysis of eluates from the DarB pulldown experiment with *Bacillus subtilis* cell extract**

Peptides identified by search of MS/MS2 data against *B. subtilis* specific protein database (UniProt Proteome ID UP000001570)

| **LK09-12: Negative control (empty vector without DarB)** | | | | |
| --- | --- | --- | --- | --- |
| **Accession** | **Description** | **Coverage (%)** | **Number of identified peptides** | **Number of peptide sequence matches** |
| P35163 | ResD | 45 | 12 | 87 |
| O07603 | YhfE | 38 | 11 | 46 |
| P27206 | SrfAA | 11 | 27 | 43 |
| P33166 | Tuf | 40 | 11 | 38 |
| Q04747 | SrfAB | 8 | 20 | 32 |
| O07937 | YraK | 22 | 5 | 29 |
| P02968 | Hag | 37 | 7 | 27 |
| P51833 | Rnc | 29 | 6 | 26 |
| O34425 | GapB | 30 | 8 | 25 |
| P46352 | XerD | 26 | 7 | 18 |
| O31986 | SunS | 23 | 8 | 17 |
| P94461 | PriA | 16 | 10 | 15 |
| P39776 | XerC | 29 | 6 | 14 |
| P42175 | NarG | 10 | 9 | 13 |
| O34803 | YeaA | 19 | 5 | 13 |
| P50735 | GudB | 12 | 4 | 12 |
| P42919 | RplB | 18 | 4 | 11 |
| P19466 | MtrB | 56 | 3 | 11 |
| P54956 | YxeQ | 20 | 6 | 10 |
| P71073 | AdeR | 15 | 5 | 10 |
| P40778 | MurC | 21 | 7 | 10 |
| P36949 | RbsB | 21 | 5 | 9 |
| P37871 | RpoC | 9 | 7 | 9 |
| P25994 | PyrAB | 11 | 8 | 9 |
| P16971 | RecA | 23 | 5 | 7 |
| Q01465 | MreB | 17 | 4 | 6 |
| O34433 | YobO | 8 | 4 | 6 |
| P54530 | YqiS | 23 | 4 | 6 |
| O31698 | DarB | 27 | 3 | 6 |
| O31475 | YcgT | 15 | 3 | 6 |
| O32158 | YurQ | 28 | 3 | 6 |
| O31774 | Rny | 4 | 2 | 6 |
| P06535 | Spo0B | 21 | 3 | 5 |
| P38021 | RocD | 7 | 2 | 5 |
| P13799 | DegS | 11 | 4 | 5 |
| P94536 | YsfB | 12 | 3 | 5 |
| P54466 | YqfA | 18 | 4 | 5 |
| P21465 | RpsC | 21 | 3 | 5 |
| P24327 | PrsA | 9 | 2 | 4 |
| O34934 | NadK2 | 11 | 2 | 4 |
| P42296 | YxiD | 7 | 3 | 4 |
| P19669 | Tal | 21 | 3 | 4 |
| P18256 | ThrZ | 6 | 3 | 4 |
| P80886 | SucC | 12 | 4 | 4 |
| P50728 | YpbB | 7 | 2 | 4 |
| O31723 | YlmA | 17 | 3 | 4 |
| P21471 | RpsJ | 43 | 3 | 4 |
| P94356 | YxkC | 22 | 2 | 3 |
| P21473 | RpsO | 10 | 2 | 3 |
| P36947 | RbsA | 5 | 2 | 3 |
| O06744 | YitI | 7 | 1 | 3 |
| P24141 | OppA | 4 | 2 | 3 |
| P94535 | GlcD | 11 | 3 | 3 |
| P50830 | YprA | 6 | 3 | 3 |
| O34384 | YceE | 15 | 2 | 3 |
| P21466 | RpsD | 19 | 2 | 3 |
| O07521 | YhaM | 4 | 1 | 3 |
| O31689 | YkvY | 9 | 3 | 3 |
| Q03523 | MurE | 9 | 3 | 3 |
| O32164 | SufS | 11 | 3 | 3 |
| P18157 | GlpK | 8 | 3 | 3 |
| O34529 | PfkA | 12 | 2 | 3 |
| P37105 | Ffh | 4 | 1 | 3 |
| P04969 | RpsK | 21 | 2 | 2 |
| P39066 | AcuB | 7 | 1 | 2 |
| P39846 | PpsB | 1 | 2 | 2 |
| P94459 | PpsD | 1 | 2 | 2 |
| P80861 | YjlD | 9 | 2 | 2 |
| P42920 | RplC | 15 | 2 | 2 |
| O34305 | YtoQ | 18 | 2 | 2 |
| P54420 | AsnB | 2 | 1 | 2 |
| P46337 | IolR | 8 | 1 | 2 |
| P39138 | RocF | 10 | 2 | 2 |
| P54563 | YqjZ | 29 | 2 | 2 |
| P40924 | Pgk | 9 | 2 | 2 |
| Q05852 | GtaB | 10 | 2 | 2 |
| P94534 | GlcF | 6 | 2 | 2 |
| O34983 | YoaP | 5 | 2 | 2 |
| O34788 | BdhA | 5 | 1 | 2 |
| P37522 | Soj | 8 | 1 | 2 |
| P12875 | RplN | 16 | 2 | 2 |
| P08065 | SdhA | 4 | 2 | 2 |
| P37870 | RpoB | 2 | 2 | 2 |
| O31645 | ManP | 4 | 2 | 2 |
| P0CI73 | GlmS | 2 | 1 | 2 |
| P17620 | RibBA | 6 | 2 | 2 |
| P21464 | RpsB | 5 | 1 | 2 |
| P19582 | Hom | 7 | 2 | 2 |
| O06478 | YfmT | 6 | 2 | 2 |
| P05654 | PyrB | 10 | 2 | 2 |
| C0SPB8 | YvaV | 15 | 2 | 2 |
| P39847 | PpsC | 1 | 2 | 2 |
| P54470 | YqfL | 11 | 2 | 2 |
| O31777 | Kbl | 8 | 2 | 2 |
| P21883 | PdhC | 11 | 2 | 2 |
| O34385 | MntA | 8 | 2 | 2 |
| P37809 | AtpD | 9 | 2 | 2 |
| Q04795 | DapG | 4 | 1 | 2 |
| P35136 | SerA | 5 | 2 | 2 |
| Q08787 | SrfAC | 1 | 1 | 1 |
| P71088 | Spo0M | 7 | 1 | 1 |
| P39634 | RocA | 3 | 1 | 1 |
| O32047 | SecDF | 2 | 1 | 1 |
| P20277 | RplQ | 13 | 1 | 1 |
| P40737 | YxxD | 12 | 1 | 1 |
| P16263 | OdhB | 4 | 1 | 1 |
| P28598 | GroL | 2 | 1 | 1 |
| P05649 | DnaN | 4 | 1 | 1 |
| O32117 | YutJ | 3 | 1 | 1 |
| O34324 | AcoL | 2 | 1 | 1 |
| O31611 | YjbM | 6 | 1 | 1 |
| O05494 | YdhC | 6 | 1 | 1 |
| P54419 | MetK | 3 | 1 | 1 |
| P12877 | RplE | 8 | 1 | 1 |
| P25996 | PyrD | 6 | 1 | 1 |
| P45913 | YqaP | 3 | 1 | 1 |
| O31664 | MtnU | 5 | 1 | 1 |
| P46354 | PunA | 7 | 1 | 1 |
| P54722 | YfiF | 5 | 1 | 1 |
| O31749 | PyrH | 5 | 1 | 1 |
| P94421 | YclQ | 4 | 1 | 1 |
| O34525 | SppA | 5 | 1 | 1 |
| O34863 | UvrA | 1 | 1 | 1 |
| P30950 | HemB | 4 | 1 | 1 |
| O31742 | RplS | 8 | 1 | 1 |
| P80871 | YwrO | 8 | 1 | 1 |
| P39751 | Mbl | 5 | 1 | 1 |
| Q03224 | GlpX | 6 | 1 | 1 |
| P94359 | YxkF | 5 | 1 | 1 |
| O34628 | YvlB | 5 | 1 | 1 |
| O34666 | CtpA | 3 | 1 | 1 |
| P37808 | AtpA | 4 | 1 | 1 |
| P08877 | PtsH | 14 | 1 | 1 |
| O07578 | YhdI | 3 | 1 | 1 |
| O34632 | SirB | 8 | 1 | 1 |
| P46899 | RplR | 13 | 1 | 1 |
| P37580 | FhuD | 5 | 1 | 1 |
| P21472 | RpsL | 8 | 1 | 1 |
| O34860 | RsbRB | 5 | 1 | 1 |
| O34797 | CoaD | 10 | 1 | 1 |

| **LK13-16: DarB coupled to Ni^2+-^NTA column** | | | | |
| --- | --- | --- | --- | --- |
| **Accession** | **Description** | **Coverage (%)** | **Number of identified peptides** | **Number of peptide sequence matches** |
| O31698 | DarB | 95 | 9 | 649 |
| O07603 | YhfE | 57 | 16 | 147 |
| O54408 | Rel | 50 | 32 | 134 |
| P33166 | TufA | 80 | 22 | 127 |
| Q04747 | SrfAB | 31 | 79 | 120 |
| P27206 | SrfAA | 29 | 70 | 113 |
| O34425 | GapB | 76 | 19 | 70 |
| P25994 | PyrAB | 39 | 31 | 57 |
| P42175 | NarG | 31 | 27 | 56 |
| P94461 | PriA | 32 | 23 | 46 |
| P80886 | SucC | 62 | 19 | 37 |
| P54420 | AsnB | 43 | 20 | 36 |
| P37871 | RpoC | 32 | 24 | 35 |
| P42920 | RplC | 31 | 6 | 27 |
| P35163 | ResD | 44 | 9 | 26 |
| P80868 | FusA | 35 | 15 | 26 |
| P46320 | LicH | 51 | 17 | 25 |
| P16971 | RecA | 49 | 10 | 25 |
| P37870 | RpoB | 19 | 15 | 25 |
| P19466 | MtrB | 75 | 6 | 24 |
| O34788 | BdhA | 46 | 10 | 23 |
| Q9KWU4 | Pyc | 22 | 17 | 22 |
| P50735 | GudB | 32 | 10 | 21 |
| P18158 | GlpD | 33 | 14 | 21 |
| P18157 | GlpK | 33 | 12 | 21 |
| P51833 | RnC | 23 | 5 | 21 |
| P37869 | Eno | 51 | 15 | 21 |
| P21467 | RpsE | 66 | 10 | 20 |
| P21465 | RpsC | 30 | 6 | 19 |
| P09339 | CitB | 20 | 13 | 19 |
| P12877 | RplE | 47 | 7 | 18 |
| P71073 | AdeR | 38 | 13 | 18 |
| P02968 | Hag | 53 | 8 | 16 |
| P42919 | RplB | 16 | 4 | 16 |
| P24141 | OppA | 31 | 12 | 16 |
| O34803 | YeaA | 37 | 8 | 16 |
| P28598 | GroL | 34 | 13 | 16 |
| P45745 | DhbF | 7 | 12 | 16 |
| P39793 | PonA | 19 | 11 | 16 |
| P37571 | ClpC/MecB | 23 | 12 | 16 |
| P70974 | RplM | 54 | 7 | 15 |
| P16263 | OdhB | 38 | 9 | 14 |
| Q01465 | MreB | 30 | 7 | 14 |
| O07937 | YraK | 25 | 5 | 14 |
| O31986 | SunS | 30 | 9 | 14 |
| P21471 | RpsJ | 44 | 4 | 14 |
| P13242 | PyrG | 25 | 10 | 14 |
| P17820 | DnaK | 13 | 5 | 14 |
| P40924 | PgK | 39 | 10 | 13 |
| P21469 | RpsG | 44 | 6 | 13 |
| P50849 | Pnp | 20 | 11 | 13 |
| O31774 | Rny | 16 | 8 | 13 |
| P04969 | RpsK | 31 | 4 | 12 |
| P08065 | SdhA | 25 | 10 | 12 |
| P14577 | RplP | 19 | 2 | 12 |
| O32222 | CsoR | 63 | 5 | 12 |
| P21476 | RpsS | 43 | 3 | 12 |
| P94459 | PpsD | 5 | 11 | 12 |
| P50830 | YprA | 17 | 10 | 12 |
| P42921 | RplD | 28 | 4 | 12 |
| P39596 | EfeM | 29 | 8 | 11 |
| P36947 | RbsA | 23 | 8 | 11 |
| O34705 | YtpA | 28 | 5 | 11 |
| P37809 | AtpD | 27 | 9 | 11 |
| O31777 | KbI | 29 | 8 | 10 |
| O34628 | YvlB | 27 | 7 | 10 |
| P21464 | RpsB | 48 | 8 | 10 |
| P42971 | PbpC | 15 | 8 | 10 |
| P23129 | OdhA | 10 | 6 | 10 |
| O31689 | YkvY | 23 | 6 | 10 |
| P38021 | RocD | 25 | 8 | 10 |
| Q7WY72 | YlzA | 57 | 5 | 10 |
| O32164 | SufS | 31 | 8 | 10 |
| P21879 | GuaB | 23 | 6 | 10 |
| Q06797 | RplA | 29 | 5 | 10 |
| P17631 | DnaJ | 23 | 6 | 10 |
| P80698 | Tig | 26 | 7 | 10 |
| O06478 | YfmT | 24 | 8 | 9 |
| P30949 | HemL | 20 | 5 | 9 |
| P39845 | PpsA | 5 | 8 | 9 |
| O32158 | YurQ | 40 | 4 | 9 |
| Q03224 | GlpX | 23 | 5 | 9 |
| O34863 | UvrA | 11 | 8 | 9 |
| Q03222 | Rho | 25 | 8 | 9 |
| O31753 | DxR | 28 | 8 | 9 |
| P13799 | DegS | 20 | 6 | 9 |
| P37570 | McsB | 33 | 8 | 9 |
| O34433 | YobO | 14 | 7 | 9 |
| P21880 | PdhD | 19 | 8 | 9 |
| Q08787 | SrfAC | 12 | 7 | 9 |
| P21473 | RpsO | 42 | 3 | 9 |
| P39847 | PpsC | 5 | 9 | 9 |
| P39846 | PpsB | 4 | 7 | 8 |
| P50866 | ClpX | 17 | 6 | 8 |
| P71011 | AlbA | 14 | 5 | 8 |
| P28264 | FtsA | 30 | 7 | 8 |
| P94547 | LcfA | 18 | 7 | 8 |
| P17620 | RibBA | 25 | 8 | 8 |
| P39778 | ClpY | 15 | 5 | 8 |
| Q05852 | GtaB | 22 | 5 | 8 |
| O34324 | AcoL | 19 | 7 | 8 |
| P17889 | InfB | 16 | 7 | 8 |
| P09124 | GapA | 27 | 6 | 8 |
| P35136 | SerA | 18 | 6 | 8 |
| P80860 | Pgi | 22 | 6 | 7 |
| O31742 | RplS | 27 | 4 | 7 |
| P54418 | PckA | 11 | 4 | 7 |
| O31541 | YetL | 32 | 5 | 7 |
| P94536 | YsfB | 23 | 6 | 7 |
| P21466 | RpsD | 37 | 5 | 7 |
| P40778 | MurC | 15 | 5 | 7 |
| Q796K8 | PbpH | 12 | 5 | 7 |
| P54956 | YxeQ | 17 | 5 | 7 |
| O34334 | YjoA | 51 | 6 | 7 |
| P19582 | Hom | 24 | 7 | 7 |
| P37877 | AckA | 21 | 6 | 7 |
| P21881 | PdhA | 28 | 7 | 7 |
| P21883 | PdhC | 24 | 7 | 7 |
| P96614 | CshA | 20 | 7 | 7 |
| P21472 | RpsL | 28 | 2 | 6 |
| P21470 | RpsI | 44 | 3 | 6 |
| P37538 | YaaQ | 25 | 2 | 6 |
| O34303 | YdjA | 13 | 6 | 6 |
| P50728 | YpbB | 12 | 3 | 6 |
| P94534 | GlcF | 17 | 6 | 6 |
| P05652 | GyrB | 8 | 3 | 6 |
| P23914 | LevR | 7 | 5 | 6 |
| P26908 | RplU | 47 | 3 | 6 |
| P80700 | tsf | 17 | 3 | 6 |
| O06491 | GatA | 17 | 5 | 6 |
| P42296 | YxiD | 12 | 5 | 6 |
| P21468 | RpsF | 48 | 3 | 6 |
| O34687 | RpmF | 27 | 1 | 6 |
| P17922 | PheT | 8 | 4 | 6 |
| P45694 | Tkt | 10 | 4 | 6 |
| P0CI73 | GlmS | 15 | 6 | 6 |
| P08838 | PtsI | 12 | 5 | 6 |
| P42060 | RplV | 56 | 4 | 6 |
| O34633 | YjlC | 37 | 4 | 6 |
| P39126 | Icd | 15 | 5 | 5 |
| Q08352 | Ald | 24 | 5 | 5 |
| P42923 | RplJ | 18 | 4 | 5 |
| P12873 | RpmC | 35 | 2 | 5 |
| P46898 | RplF | 28 | 4 | 5 |
| P50863 | SalA | 17 | 4 | 5 |
| P20429 | RpoA | 15 | 5 | 5 |
| P37465 | MetG | 8 | 4 | 5 |
| P42297 | YxiE | 39 | 4 | 5 |
| P31103 | Ndk | 34 | 4 | 5 |
| P24327 | PrsA | 9 | 2 | 5 |
| Q795M6 | YugH | 19 | 5 | 5 |
| P40806 | PksJ | 1 | 5 | 5 |
| P54419 | MetK | 13 | 4 | 5 |
| P81100 | YceC | 25 | 4 | 5 |
| P37808 | AtpA | 12 | 4 | 5 |
| Q06796 | RplK | 30 | 3 | 5 |
| O35033 | CoaBC | 16 | 5 | 5 |
| P12425 | GlnA | 20 | 5 | 5 |
| O34660 | DhaS | 11 | 4 | 5 |
| P02394 | RplL | 34 | 4 | 5 |
| O34580 | PcrA | 7 | 4 | 4 |
| P23479 | SbcD | 12 | 3 | 4 |
| Q03523 | MurE | 10 | 3 | 4 |
| O34714 | OxdC | 14 | 4 | 4 |
| P37814 | AtpF | 20 | 3 | 4 |
| P17865 | FtsZ | 15 | 4 | 4 |
| P39814 | TopA | 8 | 4 | 4 |
| P80864 | Tpx | 37 | 4 | 4 |
| O32165 | SufD | 11 | 4 | 4 |
| P37474 | Mfd | 5 | 4 | 4 |
| P39912 | AroA | 14 | 4 | 4 |
| O31760 | RnjB | 11 | 4 | 4 |
| P80885 | Pyk | 12 | 4 | 4 |
| P18256 | ThrZ | 8 | 4 | 4 |
| P49814 | Mdh | 17 | 3 | 4 |
| O34529 | PfkA | 9 | 2 | 4 |
| O31761 | YmfC | 16 | 3 | 4 |
| P29072 | CheA | 6 | 3 | 4 |
| P80865 | SucD | 11 | 2 | 4 |
| P36949 | RbsB | 11 | 3 | 4 |
| P54574 | Fur | 27 | 3 | 4 |
| P37525 | YaaB | 40 | 2 | 4 |
| P39148 | GlyA | 12 | 4 | 4 |
| O34526 | AlaS | 7 | 4 | 4 |
| P71086 | PerR | 18 | 2 | 4 |
| P05649 | DnaN | 11 | 3 | 4 |
| P54377 | GcvPB | 7 | 2 | 4 |
| O32117 | YutJ | 12 | 3 | 3 |
| P36430 | LeuS | 6 | 3 | 3 |
| O07631 | TypA | 6 | 3 | 3 |
| P37556 | YabN | 10 | 3 | 3 |
| O32090 | PncB | 6 | 2 | 3 |
| P94360 | MsmX | 12 | 3 | 3 |
| P94545 | MutSB | 4 | 2 | 3 |
| P12874 | RpsQ | 13 | 2 | 3 |
| P37477 | LysS | 8 | 3 | 3 |
| O34305 | YtoQ | 26 | 3 | 3 |
| P54523 | Dxs | 6 | 3 | 3 |
| P19946 | RplO | 35 | 3 | 3 |
| O34885 | YdiS | 7 | 2 | 3 |
| O34340 | FabF | 8 | 2 | 3 |
| P39214 | McpA | 7 | 3 | 3 |
| O31646 | ManA | 8 | 2 | 3 |
| P45913 | YqaP | 10 | 3 | 3 |
| P39633 | RocG | 9 | 3 | 3 |
| P54547 | Zwf | 9 | 3 | 3 |
| P54170 | YphP | 18 | 2 | 3 |
| P12879 | RpsH | 22 | 2 | 3 |
| O31545 | YfjO | 8 | 2 | 3 |
| O32038 | AspS | 7 | 3 | 3 |
| P37494 | YybJ | 13 | 2 | 3 |
| P08821 | HupA | 28 | 2 | 3 |
| O31626 | YjcD | 7 | 3 | 3 |
| P80870 | YugI | 33 | 3 | 3 |
| P08877 | PtsH | 42 | 2 | 3 |
| P80859 | GndA | 5 | 2 | 3 |
| P71067 | LutP | 3 | 1 | 3 |
| P28366 | SecA | 4 | 3 | 3 |
| O31749 | PyrH | 14 | 2 | 3 |
| P54530 | YqiS | 17 | 3 | 3 |
| O30509 | GatB | 9 | 3 | 3 |
| P94527 | EgsA | 11 | 3 | 3 |
| P38494 | YpfD | 11 | 3 | 3 |
| P13484 | TagE | 6 | 3 | 3 |
| P42430 | YkyB | 23 | 3 | 3 |
| O32162 | SufB | 8 | 3 | 3 |
| Q45493 | RnjA | 8 | 3 | 3 |
| P53001 | AspB | 7 | 2 | 2 |
| P55872 | InfC | 14 | 2 | 2 |
| P12042 | PurL | 4 | 2 | 2 |
| O31534 | HmoA | 19 | 2 | 2 |
| P39760 | KtrC | 10 | 2 | 2 |
| O31605 | YjbG | 4 | 2 | 2 |
| P39772 | AsnS | 5 | 1 | 2 |
| P28368 | YvyD | 6 | 1 | 2 |
| P39120 | CitZ | 7 | 2 | 2 |
| P39812 | GltA | 2 | 2 | 2 |
| P39773 | GpmI | 5 | 2 | 2 |
| O32176 | FadE | 5 | 2 | 2 |
| O31849 | YojO | 2 | 1 | 2 |
| P54518 | YqhT | 10 | 2 | 2 |
| P55873 | RplT | 24 | 2 | 2 |
| O31672 | MhqR | 17 | 2 | 2 |
| P23478 | AddA | 2 | 2 | 2 |
| P12045 | PurK | 9 | 2 | 2 |
| P07343 | FumC | 5 | 2 | 2 |
| Q05873 | ValS | 3 | 2 | 2 |
| P37942 | BfmBB | 4 | 1 | 2 |
| P54159 | YpbR | 2 | 2 | 2 |
| O31497 | YezC | 19 | 2 | 2 |
| O08455 | YhaN | 3 | 2 | 2 |
| P29727 | GuaA | 5 | 2 | 2 |
| Q45598 | YydD | 4 | 2 | 2 |
| O34996 | PolA | 3 | 2 | 2 |
| O31784 | PksR | 1 | 2 | 2 |
| O05494 | YdhC | 6 | 1 | 2 |
| P71018 | PlsX | 8 | 2 | 2 |
| P26901 | KatA | 6 | 2 | 2 |
| P28599 | GroS | 23 | 2 | 2 |
| P12875 | RplN | 14 | 1 | 2 |
| P71021 | DivIVA | 18 | 2 | 2 |
| O34591 | AcoB | 9 | 2 | 2 |
| P46318 | LicB | 11 | 1 | 2 |
| P54394 | DinG | 3 | 2 | 2 |
| P39123 | GlgP | 3 | 2 | 2 |
| P42294 | YxiB | 10 | 1 | 2 |
| P37105 | Ffh | 6 | 2 | 2 |
| P37949 | LepA | 3 | 1 | 2 |
| P54512 | MntR | 22 | 2 | 2 |
| P94542 | ZapA | 27 | 2 | 2 |
| Q45066 | ParC | 4 | 2 | 2 |
| P54375 | SodA | 8 | 1 | 2 |
| P20282 | RpsM | 16 | 2 | 2 |
| O05405 | YrhO | 7 | 1 | 2 |
| P39067 | AcuC | 4 | 1 | 2 |
| P39794 | TreP | 3 | 1 | 2 |
| P80643 | AcpA | 19 | 1 | 2 |
| P37945 | lon1 | 4 | 2 | 2 |
| P54464 | YqeY | 20 | 2 | 2 |
| P49851 | YkhA | 9 | 1 | 2 |
| O06744 | YitI | 17 | 2 | 2 |
| P21474 | RpsP | 21 | 2 | 2 |
| P54381 | GlyS | 4 | 2 | 2 |
| O34949 | YkoM | 21 | 2 | 2 |
| P46352 | XerD | 10 | 2 | 2 |
| O34348 | YfmC | 10 | 2 | 2 |
| P94523 | AraA | 6 | 2 | 2 |
| P80876 | YfkM | 15 | 1 | 2 |
| P80244 | ClpP | 7 | 1 | 2 |
| P32081 | CspB | 25 | 1 | 2 |
| P46899 | RplR | 20 | 2 | 2 |
| P20277 | RplQ | 13 | 1 | 2 |
| P49850 | MutL | 6 | 2 | 2 |
| Q04795 | DapG | 7 | 2 | 2 |
| P54542 | YqjE | 7 | 2 | 2 |
| P39776 | XerC | 10 | 2 | 2 |
| O31498 | LigA | 2 | 1 | 1 |
| P42295 | YxiC | 13 | 1 | 1 |
| P24281 | YaaK | 13 | 1 | 1 |
| O34962 | YtsJ | 5 | 1 | 1 |
| P39215 | McpB | 3 | 1 | 1 |
| P08874 | AbrB | 11 | 1 | 1 |
| P94391 | PutC | 3 | 1 | 1 |
| P12013 | GntZ | 3 | 1 | 1 |
| P46906 | ArgS | 2 | 1 | 1 |
| Q00828 | RapA | 2 | 1 | 1 |
| P54454 | YqeI | 16 | 1 | 1 |
| P37518 | YchF | 4 | 1 | 1 |
| Q7BVT7 | YerC | 13 | 1 | 1 |
| P21882 | PdhB | 4 | 1 | 1 |
| P37455 | SsbA | 10 | 1 | 1 |
| O06714 | SbcCD | 1 | 1 | 1 |
| O31766 | YmfH | 3 | 1 | 1 |
| O31661 | KinE | 1 | 1 | 1 |
| O31796 | Hfq | 14 | 1 | 1 |
| P77837 | UreC | 2 | 1 | 1 |
| P13243 | FbaA | 5 | 1 | 1 |
| O07624 | YwiB | 11 | 1 | 1 |
| O05514 | ThiL | 5 | 1 | 1 |
| P25995 | PyrC | 3 | 1 | 1 |
| P71006 | AlbF | 3 | 1 | 1 |
| P54534 | YqiW | 8 | 1 | 1 |
| P25993 | PyrAA | 3 | 1 | 1 |
| P36946 | RbsD | 12 | 1 | 1 |
| O34894 | EzrA | 2 | 1 | 1 |
| P70965 | MurAA | 4 | 1 | 1 |
| Q07428 | NrgB | 9 | 1 | 1 |
| O31575 | RecX | 3 | 1 | 1 |
| O07614 | YhfO | 10 | 1 | 1 |
| P18255 | ThrS | 2 | 1 | 1 |
| P80871 | YwrO | 8 | 1 | 1 |
| O31718 | YkzG | 17 | 1 | 1 |
| P54470 | YqfL | 4 | 1 | 1 |
| O32233 | SecG | 24 | 1 | 1 |
| O31489 | YdcI | 2 | 1 | 1 |
| Q59HN7 | PhrH | 56 | 1 | 1 |
| P54480 | YqfW | 11 | 1 | 1 |
| P56849 | RpmGA | 27 | 1 | 1 |
| Q04797 | Asd | 5 | 1 | 1 |
| P49787 | AccC | 3 | 1 | 1 |
| P46912 | GcrB | 8 | 1 | 1 |
| P39813 | DprA | 5 | 1 | 1 |
| P19947 | RpmD | 24 | 1 | 1 |
| O32178 | FadN | 2 | 1 | 1 |
| P36945 | RbsK | 6 | 1 | 1 |
| O07009 | CycB | 4 | 1 | 1 |
| P71007 | AlbE | 4 | 1 | 1 |
| P05653 | GyrA | 1 | 1 | 1 |
| O32177 | FadA | 4 | 1 | 1 |
| P25996 | PyrD | 6 | 1 | 1 |
| Q45597 | Fbp | 2 | 1 | 1 |
| O34594 | CccB | 14 | 1 | 1 |
| P06224 | SigA | 3 | 1 | 1 |
| P54423 | WprA | 2 | 1 | 1 |
| O34612 | YjlB | 8 | 1 | 1 |
| P20691 | AroA | 4 | 1 | 1 |
| O31475 | YcgT | 4 | 1 | 1 |
| P45693 | SpoVS | 17 | 1 | 1 |
| P20166 | PtsG | 3 | 1 | 1 |
| Q45494 | YkrA | 5 | 1 | 1 |
| P39586 | YwbC | 11 | 1 | 1 |
| P80861 | YjlD | 4 | 1 | 1 |
| O34481 | YrrC | 2 | 1 | 1 |
| O31648 | YjdG | 8 | 1 | 1 |
| P37572 | RadA | 3 | 1 | 1 |
| P80866 | SufC | 7 | 1 | 1 |
| C0SPA7 | YukB | 1 | 1 | 1 |
| P39066 | AcuB | 7 | 1 | 1 |
| P24012 | CtaE | 6 | 1 | 1 |
| O34667 | LuxS | 9 | 1 | 1 |
| O34629 | YerH | 4 | 1 | 1 |
| P37580 | FhuD | 5 | 1 | 1 |
| O34557 | Rpe | 8 | 1 | 1 |

**Supplementary Table 2**

**Analysis of excised gel bands from the *in vitro* pulldown experiment with Strep-Rel**

Peptides identified by search of MS/MS2 data against *B. subtilis* specific protein database (UniProt Proteome ID UP000001570)

**List of the samples**

LK29	Excised gel band 1 (Rel, DarB)

LK30	Excised gel band 2 (Rel, DarB, c-di-AMP)

LK31	Excised gel band 3 (Rel^NTD^, DarB)

LK32	Excised gel band 4 (Rel^NTD^), DarB, c-di-AMP)

| **LK29** | | | | |
| --- | --- | --- | --- | --- |
| **Accession** | **Description** | **Coverage (%)** | **Number of identified peptides** | **Number of peptide sequence matches** |
| O31698 | DarB | 89 | 8 | 576 |
| O54408 | Rel | 24 | 14 | 140 |

| **LK30** | | | | |
| --- | --- | --- | --- | --- |
| **Accession** | **Description** | **Coverage (%)** | **Number of identified peptides** | **Number of peptide sequence matches** |
| O54408 | Rel | 41 | 30 | 234 |
| O31698 | DarB | 89 | 8 | 164 |
| Q06796 | RplK | 6 | 1 | 2 |
| P12877 | RplE | 7 | 1 | 2 |

| **LK31** | | | | |
| --- | --- | --- | --- | --- |
| **Accession** | **Description** | **Coverage (%)** | **Number of identified peptides** | **Number of peptide sequence matches** |
| O31698 | DarB | 89 | 11 | 1659 |
| O54408 | Rel | 24 | 13 | 193 |
| Q06796 | RplK | 6 | 1 | 3 |

| **LK32** | | | | |
| --- | --- | --- | --- | --- |
| **Accession** | **Description** | **Coverage (%)** | **Number of identified peptides** | **Number of peptide sequence matches** |
| O31698 | DarB | 89 | 8 | 212 |
| O54408 | Rel | 29 | 20 | 118 |
| P18156 | GlpF | 10 | 1 | 1 |
| P54469 | YqfD | 2 | 1 | 1 |

**Supplementary Table 3**

**Analysis of the *in vivo* interaction experiment of DarB-Strep with low (0.1 mM) and high (5 mM) potassium concentrations**

Peptides identified by search of MS/MS2 data against *B. subtilis* specific protein database (UniProt Proteome ID UP000001570)

**List of the samples**

LK25	0.1 mM KCl, empty vector control, last wash fraction

LK26	0.1 mM KCl, empty vector control, elution fraction

LK27	0.1 mM KCl, DarB-Strep, last wash fraction

LK28	0.1 mM KCl, DarB-Strep, elution fraction

LK33	5 mM KCl, empty vector control, last wash fraction

LK34	5 mM KCl, empty vector control, elution fraction

LK35	5 mM KCl, DarB-Strep, last wash fraction

LK36	5 mM KCl, DarB-Strep, elution fraction

| **LK25** | | | | |
| --- | --- | --- | --- | --- |
| **Accession** | **Description** | **Coverage (%)** | **Number of identified peptides** | **Number of peptide sequence matches** |
| O34313 | YfkN | 1 | 1 | 5 |
| O34674 | MurJ | 3 | 1 | 1 |
| O31576 | YfhH | 17 | 1 | 1 |
| 16078234 | ThiG | 3 | 1 | 1 |
| O34443 | Apt | 4 | 1 | 1 |

| **LK26** | | | | |
| --- | --- | --- | --- | --- |
| **Accession** | **Description** | **Coverage (%)** | **Number of identified peptides** | **Number of peptide sequence matches** |
| Q9KWU4 | Pyc | 63 | 57 | 377 |
| P28598 | GroL | 30 | 12 | 26 |
| P19582 | Hom | 29 | 11 | 23 |
| P02968 | Hag | 37 | 7 | 21 |
| P33166 | Tuf | 26 | 8 | 20 |
| P17820 | DnaK | 24 | 10 | 17 |
| P40924 | Pgk | 26 | 8 | 17 |
| P39126 | Icd | 23 | 8 | 16 |
| P21883 | PdhC | 24 | 7 | 14 |
| P09124 | GapA | 24 | 6 | 12 |
| P16971 | RecA | 27 | 6 | 12 |
| P37808 | AtpA | 15 | 7 | 12 |
| P35136 | SerA | 14 | 6 | 11 |
| P80886 | SucC | 19 | 6 | 11 |
| P21464 | RpsB | 29 | 5 | 10 |
| P24141 | OppA | 12 | 5 | 10 |
| P08065 | SdhA | 10 | 5 | 10 |
| P37809 | AtpD | 12 | 4 | 10 |
| O54408 | Rel | 8 | 5 | 9 |
| P24327 | PrsA | 16 | 4 | 8 |
| Q05852 | GtaB | 19 | 4 | 8 |
| O32167 | MetQ | 20 | 4 | 8 |
| P39751 | mbl | 13 | 4 | 8 |
| P54535 | ArtP | 18 | 4 | 8 |
| P94565 | LeuA | 10 | 4 | 7 |
| P17631 | DnaJ | 10 | 3 | 7 |
| Q01465 | MreB | 13 | 4 | 7 |
| P37476 | FtsH | 8 | 4 | 7 |
| O07021 | LutB | 7 | 3 | 6 |
| O32157 | FrlB | 11 | 3 | 6 |
| P21465 | RpsC | 21 | 3 | 6 |
| P49786 | AccB | 33 | 3 | 6 |
| P21882 | PdhB | 9 | 3 | 6 |
| P80868 | FusA | 6 | 3 | 6 |
| O34538 | YcdA | 10 | 3 | 6 |
| P20429 | RpoA | 15 | 4 | 6 |
| P94356 | YxkC | 18 | 3 | 6 |
| O34628 | YvlB | 14 | 3 | 6 |
| P04969 | RpsK | 15 | 2 | 5 |
| P08066 | SdhB | 15 | 3 | 5 |
| P39912 | AroA | 11 | 3 | 5 |
| P25994 | PyrAB | 4 | 3 | 5 |
| P20166 | PtsG | 4 | 3 | 5 |
| P80861 | YjlD | 8 | 2 | 4 |
| O32213 | CysI | 4 | 2 | 4 |
| P37253 | IlvC | 8 | 2 | 4 |
| P50863 | SalA | 6 | 2 | 4 |
| O34358 | HtrA | 5 | 2 | 4 |
| P39765 | PyrR | 12 | 2 | 4 |
| P21879 | GuaB | 7 | 2 | 4 |
| P51835 | FtsY | 7 | 2 | 3 |
| P94421 | YclQ | 8 | 2 | 3 |
| Q06797 | RplA | 9 | 2 | 3 |
| P96740 | PgdS | 9 | 2 | 3 |
| P21472 | RpsL | 8 | 1 | 2 |
| P37814 | AtpF | 6 | 1 | 2 |
| P54466 | YqfA | 4 | 1 | 2 |
| O31749 | PyrH | 14 | 2 | 2 |
| O34788 | BdhA | 5 | 1 | 2 |
| P12045 | PurK | 5 | 1 | 2 |
| P28264 | FtsA | 2 | 1 | 2 |
| O31753 | Dxr | 3 | 1 | 2 |
| O32047 | SecDF | 2 | 1 | 2 |
| O34344 | SdpC | 5 | 1 | 2 |
| P54419 | MetK | 4 | 1 | 2 |
| P39772 | AsnS | 3 | 1 | 2 |
| O31774 | Rny | 2 | 1 | 2 |
| P46898 | RplF | 6 | 1 | 2 |
| P39778 | ClpY | 2 | 1 | 2 |
| O34347 | ArgG | 3 | 1 | 2 |
| P54576 | McpC | 3 | 1 | 2 |
| P39617 | YwdI | 16 | 1 | 2 |
| P17889 | InfB | 2 | 1 | 2 |
| P34956 | QoxB | 2 | 1 | 2 |
| P21467 | RpsE | 5 | 1 | 2 |
| P21881 | PdhA | 2 | 1 | 2 |
| P37869 | Eno | 4 | 1 | 2 |
| P37871 | RpoC | 1 | 1 | 2 |
| P54342 | XkdW | 6 | 1 | 2 |
| O34857 | Rok | 16 | 1 | 1 |
| P21880 | PdhD | 6 | 1 | 1 |
| 2632038 | MetC | 2 | 1 | 1 |
| P37105 | Ffh | 2 | 1 | 1 |
| O34992 | OpuCA | 4 | 1 | 1 |
| P37870 | RpoB | 1 | 1 | 1 |
| P40767 | CwlO | 2 | 1 | 1 |
| O31902 | YorL | 1 | 1 | 1 |
| P37527 | PdxS | 4 | 1 | 1 |
| Q03224 | GlpX | 3 | 1 | 1 |
| P35155 | ScpB | 5 | 1 | 1 |
| P37887 | CysK | 5 | 1 | 1 |

| **LK27** | | | | |
| --- | --- | --- | --- | --- |
| **Accession** | **Description** | **Coverage (%)** | **Number of identified peptides** | **Number of peptide sequence matches** |
| P02968 | Hag | 19 | 4 | 8 |
| P54518 | YqhT | 2 | 1 | 2 |
| P94584 | FabZ | 6 | 1 | 2 |
| O06714 | SbcC | 1 | 1 | 1 |
| O31489 | YdcI | 1 | 1 | 1 |
| O34520 | HisG | 6 | 1 | 1 |
| O06008 | AdhR | 9 | 1 | 1 |
| P55872 | InfC | 3 | 1 | 1 |
| P54542 | YqjE | 3 | 1 | 1 |
| O54408 | Rel | 1 | 1 | 1 |
| P46898 | RplF | 5 | 1 | 1 |
| P54453 | YqeH | 3 | 1 | 1 |
| O32101 | YueB | 1 | 1 | 1 |

| **LK28** | | | | |
| --- | --- | --- | --- | --- |
| **Accession** | **Description** | **Coverage (%)** | **Number of identified peptides** | **Number of peptide sequence matches** |
| Q9KWU4 | Pyc | 54 | 49 | 292 |
| O31698 | DarB | 88 | 8 | 187 |
| O54408 | Rel | 20 | 15 | 57 |
| P02968 | Hag | 71 | 11 | 39 |
| P33166 | Tuf | 28 | 8 | 19 |
| P21883 | PdhC | 32 | 8 | 17 |
| P21880 | PdhD | 25 | 8 | 17 |
| O32157 | FrlB | 27 | 6 | 17 |
| P28598 | GroL | 15 | 6 | 16 |
| P35136 | SerA | 17 | 7 | 14 |
| P19582 | Hom | 23 | 7 | 13 |
| P16971 | RecA | 27 | 6 | 12 |
| O34788 | BdhA | 10 | 2 | 11 |
| P21881 | PdhA | 16 | 5 | 11 |
| Q05852 | GtaB | 23 | 5 | 10 |
| P40924 | Pgk | 19 | 6 | 9 |
| P80868 | FusA | 12 | 5 | 9 |
| P37869 | Eno | 15 | 4 | 8 |
| P80886 | SucC | 15 | 4 | 8 |
| O32156 | YurO | 14 | 4 | 8 |
| O07021 | LutB | 11 | 4 | 8 |
| P39666 | NadC | 18 | 4 | 8 |
| P21882 | PdhB | 14 | 4 | 7 |
| P37809 | AtpD | 12 | 4 | 7 |
| P37808 | AtpA | 9 | 4 | 7 |
| P80861 | YjlD | 11 | 3 | 6 |
| P39126 | Icd | 9 | 3 | 6 |
| P16263 | OdhB | 10 | 3 | 6 |
| P54518 | YqhT | 10 | 2 | 6 |
| P09124 | GapA | 10 | 3 | 5 |
| P21464 | RpsB | 22 | 3 | 5 |
| P04969 | RpsK | 15 | 2 | 5 |
| P21879 | GuaB | 8 | 3 | 5 |
| O31742 | RplS | 28 | 3 | 5 |
| P08066 | SdhB | 8 | 2 | 4 |
| O32167 | MetQ | 10 | 2 | 4 |
| P24141 | OppA | 6 | 2 | 4 |
| P21467 | RpsE | 20 | 2 | 4 |
| Q01465 | MreB | 10 | 3 | 4 |
| P54535 | ArtP | 9 | 2 | 4 |
| O34628 | YvlB | 8 | 2 | 4 |
| P21465 | RpsC | 14 | 2 | 4 |
| P39912 | AroA | 5 | 1 | 4 |
| P37253 | IlvC | 8 | 2 | 4 |
| O07020 | LutA | 8 | 2 | 4 |
| P08065 | SdhA | 5 | 3 | 3 |
| P17820 | DnaK | 6 | 3 | 3 |
| P17631 | DnaJ | 7 | 2 | 3 |
| P51835 | FtsY | 6 | 2 | 3 |
| P24327 | PrsA | 8 | 2 | 3 |
| P94565 | LeuA | 5 | 2 | 3 |
| P46898 | RplF | 11 | 2 | 3 |
| P12875 | RplN | 16 | 2 | 3 |
| P49786 | AccB | 21 | 2 | 3 |
| Q01464 | MinD | 9 | 2 | 2 |
| P45740 | ThiC | 3 | 1 | 2 |
| O31749 | PyrH | 5 | 1 | 2 |
| O34538 | YcdA | 3 | 1 | 2 |
| O32213 | CysI | 2 | 1 | 2 |
| P70974 | RplM | 7 | 1 | 2 |
| P54419 | MetK | 4 | 1 | 2 |
| O31664 | MtnU | 5 | 1 | 2 |
| P39765 | PyrR | 6 | 1 | 2 |
| P42971 | PbpC | 2 | 1 | 2 |
| P80859 | GndA | 2 | 1 | 2 |
| P28264 | FtsA | 3 | 1 | 2 |
| P37105 | Ffh | 2 | 1 | 2 |
| O06478 | YfmT | 2 | 1 | 2 |
| P14949 | TrxA | 13 | 1 | 2 |
| P54466 | YqfA | 7 | 1 | 2 |
| P37476 | FtsH | 2 | 1 | 2 |
| P12877 | RplE | 12 | 1 | 2 |
| P37870 | RpoB | 1 | 1 | 2 |
| P23129 | OdhA | 1 | 1 | 2 |
| P94421 | YclQ | 4 | 1 | 2 |
| P42921 | RplD | 5 | 1 | 2 |
| O05252 | YufN | 6 | 1 | 1 |
| P37527 | PdxS | 4 | 1 | 1 |
| P54382 | FolD | 4 | 1 | 1 |
| O32259 | LutC | 6 | 1 | 1 |
| P04990 | ThrC | 4 | 1 | 1 |
| P39148 | GlyA | 3 | 1 | 1 |
| Q06797 | RplA | 3 | 1 | 1 |
| P81100 | YceC | 5 | 1 | 1 |
| P21471 | RpsJ | 7 | 1 | 1 |
| P42924 | RplW | 11 | 1 | 1 |
| P09339 | CitB | 1 | 1 | 1 |

| **LK33** | | | | |
| --- | --- | --- | --- | --- |
| **Accession** | **Description** | **Coverage (%)** | **Number of identified peptides** | **Number of peptide sequence matches** |
| O34313 | YfkN | 1 | 1 | 20 |
| O34921 | YtoI | 2 | 1 | 1 |
| P42420 | IolJ | 3 | 1 | 1 |
| P46898 | RplF | 5 | 1 | 1 |
| O34578 | YjnA | 4 | 1 | 1 |
| P54334 | XkdO | 1 | 1 | 1 |
| Q9KWU4 | Pyc | 1 | 1 | 1 |
| P54488 | YqgF | 3 | 1 | 1 |
| O31489 | YdcI | 1 | 1 | 1 |

| **LK34** | | | | |
| --- | --- | --- | --- | --- |
| **Accession** | **Description** | **Coverage (%)** | **Number of identified peptides** | **Number of peptide sequence matches** |
| Q9KWU4 | Pyc | 59 | 56 | 306 |
| P02968 | Hag | 37 | 7 | 16 |
| P49786 | AccB | 27 | 3 | 8 |
| P17820 | DnaK | 6 | 3 | 5 |
| P94356 | YxkC | 15 | 3 | 5 |
| O34857 | Rok | 6 | 2 | 4 |
| P28598 | GroL | 6 | 2 | 4 |
| O34358 | HtrA | 5 | 2 | 3 |
| P54518 | YqhT | 2 | 1 | 3 |
| O32167 | MetQ | 3 | 1 | 2 |
| P04969 | RpsK | 10 | 1 | 2 |
| P17631 | DnaJ | 3 | 1 | 2 |
| P37809 | AtpD | 3 | 1 | 2 |
| P24327 | PrsA | 4 | 1 | 2 |
| P39912 | AroA | 3 | 1 | 2 |
| P19582 | Hom | 3 | 1 | 2 |
| O34344 | SdpC | 5 | 1 | 2 |
| P14577 | RplP | 10 | 1 | 2 |
| 1644207 | RplM | 7 | 1 | 1 |
| 786159 | RpsC | 3 | 1 | 1 |
| O34921 | YtoI | 2 | 1 | 1 |
| P23129 | OdhA | 3 | 1 | 1 |
| P02394 | RplL | 7 | 1 | 1 |
| P08065 | SdhA | 2 | 1 | 1 |
| P46898 | RplF | 6 | 1 | 1 |
| C0H3Z2 | YjzH | 19 | 1 | 1 |

| **LK35** | | | | |
| --- | --- | --- | --- | --- |
| **Accession** | **Description** | **Coverage (%)** | **Number of identified peptides** | **Number of peptide sequence matches** |
| O34313 | YfkN | 1 | 1 | 32 |
| P02968 | Hag | 13 | 3 | 3 |
| Q9KWU4 | Pyc | 1 | 1 | 2 |
| P96583 | TopB | 2 | 1 | 2 |
| O34466 | YodR | 6 | 1 | 2 |
| P45693 | SpoVS | 24 | 1 | 1 |
| O34858 | ArgH | 3 | 1 | 1 |
| Q03221 | Tdk | 3 | 1 | 1 |
| O34921 | YtoI | 2 | 1 | 1 |
| O31489 | YdcI | 1 | 1 | 1 |
| O34709 | OpcR | 9 | 1 | 1 |
| O31698 | DarB | 10 | 1 | 1 |
| P54531 | YqiT | 2 | 1 | 1 |
| O06714 | SbcC | 1 | 1 | 1 |

| **LK36** | | | | |
| --- | --- | --- | --- | --- |
| **Accession** | **Description** | **Coverage (%)** | **Number of identified peptides** | **Number of peptide sequence matches** |
| O31698 | DarB | 89 | 9 | 200 |
| Q9KWU4 | Pyc | 44 | 36 | 144 |
| P02968 | Hag | 48 | 8 | 19 |
| P45740 | ThiC | 21 | 7 | 17 |
| P21883 | PdhC | 17 | 5 | 9 |
| P16263 | OdhB | 14 | 4 | 7 |
| P94565 | LeuA | 8 | 3 | 6 |
| P33166 | Tuf | 12 | 3 | 6 |
| P21882 | PdhB | 14 | 3 | 6 |
| P17820 | DnaK | 8 | 3 | 6 |
| P54518 | YqhT | 14 | 2 | 5 |
| P19582 | Hom | 6 | 2 | 4 |
| P49786 | AccB | 29 | 2 | 4 |
| P28598 | GroL | 7 | 2 | 4 |
| P16971 | RecA | 11 | 2 | 4 |
| P39666 | NadC | 16 | 3 | 4 |
| P37808 | AtpA | 6 | 2 | 4 |
| P21880 | PdhD | 8 | 2 | 3 |
| P42971 | PbpC | 5 | 2 | 3 |
| P37809 | AtpD | 6 | 2 | 3 |
| P24141 | OppA | 6 | 2 | 3 |
| P24327 | PrsA | 5 | 1 | 2 |
| P08065 | SdhA | 2 | 1 | 2 |
| P17631 | DnaJ | 4 | 1 | 2 |
| O32156 | YurO | 4 | 1 | 2 |
| P39912 | AroA | 3 | 1 | 2 |
| O07021 | LutB | 3 | 1 | 2 |
| P80643 | AcpA | 12 | 1 | 2 |
| P46898 | RplF | 6 | 1 | 2 |
| C0SPB6 | SsbB | 15 | 1 | 2 |
| O34344 | SdpC | 5 | 1 | 2 |
| O32157 | FrlB | 5 | 1 | 2 |
| P37518 | YchF | 4 | 1 | 2 |
| P37253 | IlvC | 3 | 1 | 2 |
| P14949 | TrxA | 13 | 1 | 2 |
| P80886 | SucC | 3 | 1 | 2 |
| P54342 | XkdW | 6 | 1 | 2 |
| O05252 | YufN | 6 | 1 | 2 |
| O34358 | HtrA | 4 | 1 | 1 |
| P54535 | ArtP | 5 | 1 | 1 |
| P37948 | GlpT | 4 | 1 | 1 |
| P08066 | SdhB | 5 | 1 | 1 |
| P02394 | RplL | 10 | 1 | 1 |
| O32214 | CysJ | 2 | 1 | 1 |
| O32113 | SufA | 13 | 1 | 1 |
| P53557 | BioB | 5 | 1 | 1 |
| P35155 | ScpB | 5 | 1 | 1 |
| P21465 | RpsC | 7 | 1 | 1 |
| Q01465 | MreB | 4 | 1 | 1 |
| Q45058 | CotM | 8 | 1 | 1 |
| O34347 | ArgG | 3 | 1 | 1 |
| P39846 | PpsB | 1 | 1 | 1 |

**Supplementary Table 4**

**Plasmids used in this study**

| **Name** | **Vector** | **Insert** | **Reference** |
| --- | --- | --- | --- |
| pGP635 | pGP380/ XbaI + PstI | PCR-Product *darB*, TK01/TK02 (XbaI + PstI) | This study |
| pGP706 | pWH844/ SalI | PCR-Prod. *ccpC,* HMB9/HMB11/ (SalI) | 1 |
| pGP767 | pGP382/ XbaI + PstI | PCR-Prod. *darB*, TK03/TK04 (XbaI + PstI) | This study |
| pGP2972 | pET-SUMO/ BsaI + XhoI | PCR-Prod. *darB,* LK80/LK81 (BsaI + XhoI) | This study |
| pGP2974 | pUT18/ XbaI + KpnI | PCR- Prod. *darB,* LK130/LK131 (XbaI + KpnI) | This study |
| pGP2975 | pUT18c/ XbaI + KpnI | PCR- Prod. *darB*, LK130/LK131 (XbaI + KpnI) | This study |
| pGP2976 | pKT25/ XbaI + KpnI | PCR- Prod. *darB*, LK130/LK131 *(XbaI + KpnI)* | This study |
| pGP2977 | p25-N/ XbaI + KpnI | PCR- Prod. *darB*, LK130/LK131 (XbaI + KpnI) | This study |
| pGP2982 | pUT18/ XbaI + KpnI | PCR- Prod. *ccpC,* LK161/LK162 *(XbaI + KpnI)* | This study |
| pGP2983 | pUT18c/ XbaI + KpnI | PCR- Prod. *ccpC,* LK161/LK162 (XbaI + KpnI) | This study |
| pGP2984 | pKT25/ XbaI + KpnI | PCR- Prod. *ccpC,* LK161/LK162 (XbaI + KpnI) | This study |
| pGP2985 | p25-N/ XbaI + KpnI | PCR- Prod. *ccpC,* LK161/LK162 (XbaI + KpnI) | This study |
| pGP3306 | pBQ200/ XbaI + PstI | PCR-Prod. *darB,* LK209/TK02 (XbaI + PstI) | This study |
| pGP3330 | pWH844/ BamHI + SalI | PCR-Prod. *rel,* LK283/LK284 (BamHI + SalI) | This study |
| pGP3336 | pUT18/ XbaI + KpnI | PCR- Prod. *sasA,* LK312/LK313 (XbaI + KpnI) | This study |
| pGP3337 | pUT18c/ XbaI + KpnI | PCR- Prod. *sasA,* LK312/LK313 (XbaI + KpnI) | This study |
| pGP3338 | pKT25/ XbaI + KpnI | PCR- Prod. *sasA,* LK312/LK313 (XbaI + KpnI) | This study |
| pGP3339 | p25-N/ XbaI + KpnI | PCR- Prod. *sasA,* LK312/LK313 (XbaI + KpnI) | This study |
| pGP3344 | pUT18/ XbaI + KpnI | PCR- Prod. *rel,* LK287/LK288 (XbaI + KpnI) | This study |
| pGP3345 | pUT18c/ XbaI + KpnI | PCR- Prod. *rel,* LK287/LK288 (XbaI + KpnI) | This study |
| pGP3346 | pKT25/ XbaI + KpnI | PCR- Prod. *rel,* LK287/LK288 (XbaI + KpnI) | This study |
| pGP3347 | p25-N/ XbaI + KpnI | PCR- Prod. *rel,* LK287/LK288 (XbaI + KpnI) | This study |
| pGP3348 | pGP172/ KpnI + BamHI | PCR-Prod. *rel* LK310/LK311 (KpnI + BamHI) | This study |
| pGP3350 | pGP172/ KpnI + BamHI | PCR-Produkt *rel*^NTD^ LK310/LK333 (KpnI + BamHI) | This study |
| pGP3411 | pUT18/ XbaI + KpnI | PCR- Prod. *sasB,* LK314/LK315 (XbaI + KpnI) | This study |
| pGP3412 | pUT18c/ XbaI + KpnI | PCR- Prod. *sasB,* LK314/LK315 (XbaI + KpnI) | This study |
| pGP3413 | pKT25/ XbaI + KpnI | PCR- Prod. *sasB,* LK314/LK315 (XbaI + KpnI) | This study |
| pGP3414 | p25-N/ XbaI + KpnI | PCR- Prod. *sasB,* LK314/LK315 (XbaI + KpnI) | This study |
| pGP3415 | pUT18/ XbaI + KpnI | PCR- Prod. *rel*^SYN-RRM^*,* LK316/LK288 (XbaI + KpnI) | This study |
| pGP3416 | pUT18c/ XbaI + KpnI | PCR- Prod. *rel*^SYN-RRM^*,* LK316/LK288 (XbaI + KpnI) | This study |
| pGP3417 | pKT25/ XbaI + KpnI | PCR- Prod. *rel*^SYN-RRM^*,* LK316/LK288 (XbaI + KpnI) | This study |
| pGP3418 | p25-N/XbaI + KpnI | PCR- Prod. *rel*^SYN-RRM^*,* LK316/LK288 (XbaI + KpnI) | This study |
| pGP3419 | pUT18/ XbaI + KpnI | PCR- Prod. *rel*^NTD^*,* LK287/LK317 (XbaI + KpnI) | This study |
| pGP3420 | pUT18c/ XbaI + KpnI | PCR- Prod. *rel*^NTD^, LK287/LK317 (XbaI + KpnI) | This study |
| pGP3421 | pKT25/ XbaI + KpnI | PCR- Prod. *rel*^NTD^*,* LK287/LK317 (XbaI + KpnI) | This study |
| pGP3422 | p25-N/ XbaI + KpnI | PCR- Prod. *rel*^NTD^*,* LK287/LK317 (XbaI + KpnI) | This study |
| pGP3429 | pWH844/ BamHI + SalI | PCR-Prod. *rel*^NTD^*,* LK283 + LK358 (BamHI + SalI) | This study |
| pGP3437 | pBQ200/ XbaI + PstI | PCR-Prod. *darB*^A25G^*,* LK209+ TK02 + LK372 (XbaI + PstI) | This study |
| pGP3441 | pBQ200/ XbaI + PstI | PCR-Prod. *darB*^R132M^*,* LK209+ TK02 + LK376 (XbaI + PstI) | This study |
| pGP3444 | pET-SUMO/ XhoI + BsaI | PCR-Prod. *darB*^A25G^*,* LK80 + LK81 (XhoI + BsaI) | This study |
| pGP3448 | pET-SUMO/XhoI+BsaI | PCR-Prod. *darB*^R132M^*,* LK80 + LK81 (XhoI + BsaI) | This study |
| pGP3460 | pET-SUMO/XhoI+BsaI | PCR-Prod. *darB*^A25G/R132M^*,* LK80 + LK81 (XhoI + BsaI) | This study |
| pGP3601 | pBQ200/XbaI+PstI | PCR-Prod. *darB*^A25G,R132M^*,* LK209+ TK02 + LK376 (XbaI + PstI) | This study |
| pVHP186 | pET24d | *rel* | 2 |
